# Multiple environmental signaling pathways control the differentiation of RORγt-expressing regulatory T cells

**DOI:** 10.1101/791244

**Authors:** Hind Hussein, Sébastien Denanglaire, Frédéric Van Gool, Abdulkader Azouz, Yousra Ajouaou, Hana El-Khatib, Guillaume Oldenhove, Oberdan Leo, Fabienne Andris

**Author notes:** Reprint requests and page proofs should be addressed to*: Dr. Fabienne Andris, Laboratoire d’Immunobiologie, Université Libre de Bruxelles – IBMM, 12, rue des Prof. Jeener et Brachet, 6041 Gosselies, BELGIUM.

## Abstract

RORγt-expressing Tregs form a specialized subset of intestinal CD4^+^ Foxp3^+^ cells which is essential to maintain gut homeostasis and tolerance to commensal microbiota. Recently, c-Maf emerged as a critical factor in the regulation of RORγt expression in Tregs. However, aside from c-Maf signaling, the signaling pathways involved in the differentiation of RORγt^+^ Tregs and their possible interplay with c-Maf in this process are largely unknown. We show that RORγt^+^ Treg development is controlled by positive as well as negative signals. Along with c-Maf signaling, signals derived from a complex microbiota, as well as IL-6/STAT3- and TGF-β-derived signals act in favor of RORγt^+^ Treg development. Ectopic expression of c-Maf did not rescue RORγt expression in STAT3-deficient Tregs, indicating the presence of additional effectors downstream of STAT3. Moreover, we show that an inflammatory IFN-γ/STAT1 signaling pathway acts as a negative regulator of RORγt^+^ Treg differentiation in a c-Maf independent fashion.

These data thus argue for a complex integrative signaling network that finely tunes RORγt expression in Tregs. The finding that type 1 inflammation impedes RORγt^+^ Treg development even in the presence of an active IL-6/STAT3 pathway further suggests a dominant negative effect of STAT1 over STAT3 in this process.

## Introduction

CD4 T cells expressing transcription factor forkhead box P3 (Foxp3) constitute a regulatory lineage of T cells (Treg) which maintains immune tolerance against self-antigens and prevents tissue destruction consequent to excessive immune responses. Tregs can be generated in the thymus from developing CD4^+^ thymocytes (tTregs) or can result from the differentiation of mature T cells in the periphery (pTregs) (Chen et al., 2003; Panduro et al., 2016; Whibley et al., 2019). Recent findings indicate that, similarly to conventional helper T cells, Tregs are phenotypically and functionally heterogeneous. Distinct Treg populations adopt specialized phenotypes through the co-expression of Foxp3 and lineage-defining transcription factors such as PPARγ, BCL6, or RORγt in response to tissue- or inflammatory-driven signals (Panduro et al., 2016).

In particular, RORγt^+^ Tregs are found in the intestinal tissue of naïve mice. Signals deriving from a complex microbiota as well as STAT3 signaling are necessary to their presence in the intestinal compartment (Ohnmacht et al., 2015; Sefik et al., 2015). This subset of Tregs has been shown to protect efficiently from intestinal immunopathology in different colitis models (Lochner, 2011; Ohnmacht et al., 2015; Sefik et al., 2015; Yang et al., 2015) and to mediate immunological tolerance to the gut pathobiont *Helicobacter hepaticus* (Xu et al., 2018). A recent study showed that thymic-derived Tregs also expressed RORγt in lymph nodes following immunization and were able to protect mice from Th17 cell-mediated CNS inflammation (Kim et al., 2017). Although RORγt expression can be acquired by ex-Tregs during pathogenic Th17 conversion (Komatsu, 2014), RORγt-expressing Tregs mostly represent a Treg lineage participating in the immunological tolerance of barrier tissues and protecting from autoimmunity (Kim et al., 2017; Lochner, 2011; Ohnmacht et al., 2015; Sefik et al., 2015; Xu et al., 2018; Yang et al., 2015). RORγt^+^ Tregs can also develop in the tumor microenvironment where they hinder anti-tumor immunity, thus revealing a double-edged function of this Treg subset in immune homeostasis (Downs-Canner et al., 2017). However, despite their importance in physiological and pathological immune responses, the factors driving RORγt^+^ Treg differentiation are still incompletely defined.

Recent studies reported that transcription factor c-Maf promotes the differentiation of intestinal RORγt^+^ Tregs (Imbratta et al., 2019; Wheaton et al., 2017; Xu et al., 2018). Coincidentally, transcriptomic studies conducted on Tregs originating from different tissues revealed a strong enrichment for c-Maf in the intestinal compartment (Sefik et al., 2015). Transcription factor c-Maf, a member of the AP-1 family of basic region/leucine zipper transcription factors, is expressed by distinct CD4^+^ T cell subsets, including Th17, Th2, Tfh and Tr1 cells, and is thought to regulate the expression of IL-10, IL-4, and IL-21 through the transactivation of their promoters, downstream of Batf, ICOS, and STAT3 signaling (Andris et al., 2017; Apetoh et al., 2010; Bauquet et al., 2009; Hiramatsu et al., 2010; Sahoo et al., 2015). Thus, and similarly to what has been previously described for Th17 cells (Tanaka et al., 2014a), expression of RORγt in Tregs is c-Maf-dependent. However, unlike RORγt expression, which is restricted to gut-associated Tregs in naïve mice, c-Maf is expressed by a wider proportion of Tregs found in distinct organs. Of note, high levels of c-Maf are found in a subset of splenic CD44^+^ CD62L^-^ effector Tregs driven by ICOS signaling (Wheaton et al., 2017). The partial overlapping of RORγt and c-Maf expression along with the presence of a substantial population of c-Maf^+^ RORγt^-^ Tregs in lymphoid organs therefore suggests that c-Maf is not sufficient *per se* to drive RORγt^+^ Treg cell differentiation and supports the existence of complementary signaling pathways.

Herein, extensive analysis of the lymphoid organs and tissues of genetically invalidated mice or mice harboring an altered microbiota revealed that, well beyond the c-Maf/RORγt interplay, multiple signaling pathways cooperate to exert a tight control over RORγt expression in Tregs.

## Results

### c-Maf is highly expressed in intestinal Tregs and is required for RORγt expression

We first investigated the expression of c-Maf and RORγt in Tregs found in distinct organs. In accordance with previous data (Ohnmacht et al., 2015; Sefik et al., 2015), we observed that RORγt^+^ Tregs were mainly present in the small intestine and colon lamina propria and, to a lesser extent, in mesenteric lymph nodes (Figure 1A, B). Of note, a large proportion of Tregs (ranging from 20% to 85%) expressed c-Maf in all the examined organs, with a notable exception for the thymus (Figure 1A, C). In the intestine, both RORγt^+^ and RORγt^-^ Treg subsets expressed c-Maf, although expression levels were higher in the RORγt^+^ compartment (Figure 1A, D, E). Strikingly, FACS analysis also revealed that a large fraction of c-Maf^+^ Tregs do not express RORγt. This was observed in all the examined organs and was most evident in the small intestine lamina propria, where two-thirds of the c-Maf^+^ Tregs lacked RORγt expression (Figure 1A).

**Figure 1.**
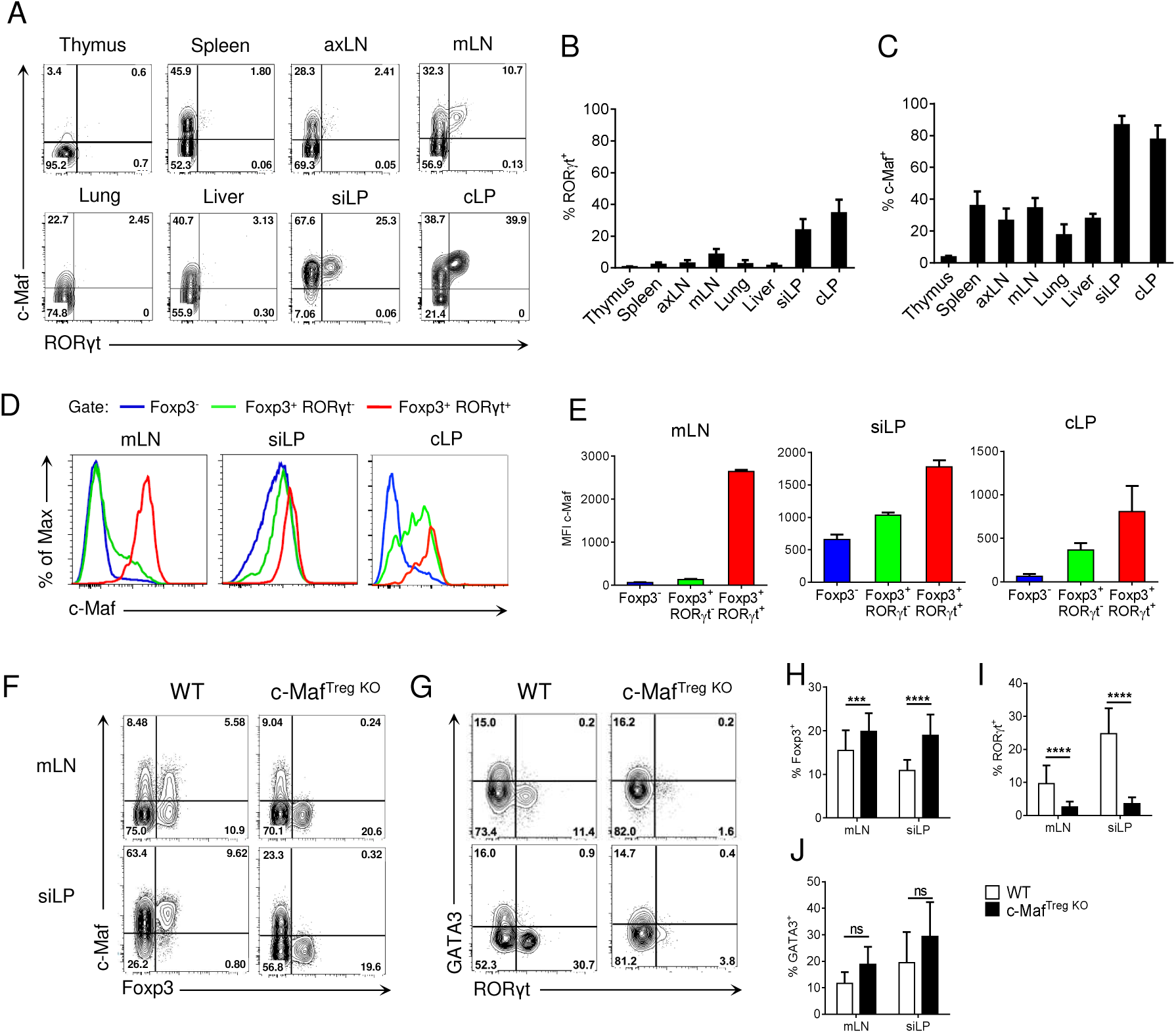
Transcription factor c-Maf is required for the differentiation of RORγt^+^ Tregs. (A) Representative flow cytometry expression profiles of c-Maf versus RORγt of Treg cells in the indicated organs of naïve C57BL/6 mice (gate CD4^+^ Foxp3^+^). (B, C) Histograms showing the frequency of RORγt^+^ (B) or c-Maf^+^ (C) Treg cells in the indicated organs. D, E) Expression profile (D) and median of fluorescence intensity (E) of c-Maf among Foxp3^-^, Foxp3^+^ RORγt^-^and Foxp3^+^ RORγt^+^ CD4 T cells in the indicated organs. (F, G) Representative flow cytometry expression profiles of c-Maf versus Foxp3 (F, gate CD4^+^) and GATA3 versus RORγt (G, gate CD4^+^ Foxp3^+^) from mLN and siLP of WT and c-Maf^Treg O^ mice. (H-J) Histograms showing the frequency of total Tregs (H), RORγt^+^ Tregs (I) and GATA3^+^ Tregs (J) in mLN and siLP of WT and c-Maf^Treg KO^ mice. Results are representative of at least three independent experiments; histograms in (B, C, E, H-J) represent the mean ± SD of at least five individual mice. Difference between groups is determined by an unpaired t test (H, I) or a Mann–Whitney test for two-tailed data (J). ***p < 0.001; ****p < 0.0001. (axLN: axillary lymph nodes, mLN: mesenteric lymph nodes, siLP: small intestine lamina propria, cLP: colon lamina propria)

In lymphoid organs, c-Maf expression was mainly found among effector Tregs, characterized by the expression of high levels of ICOS and CD44 (Figure S1 in Supplementary Material). In contrast to RORγt^+^ Tregs, which were mostly of peripheral origin, c-Maf^+^ Tregs were found both in Nrp1^+^ and Nrp1^-^ Tregs, suggesting that c-Maf^+^ Tregs can be of thymic or peripheral origin (Figure 2A, B). Thymic c-Maf^+^ Tregs were enriched in the spleen whereas their peripheral counterparts were enriched in mesenteric lymph nodes and formed the majority of intestinal Tregs (Figure 2C).

**Figure 2.**
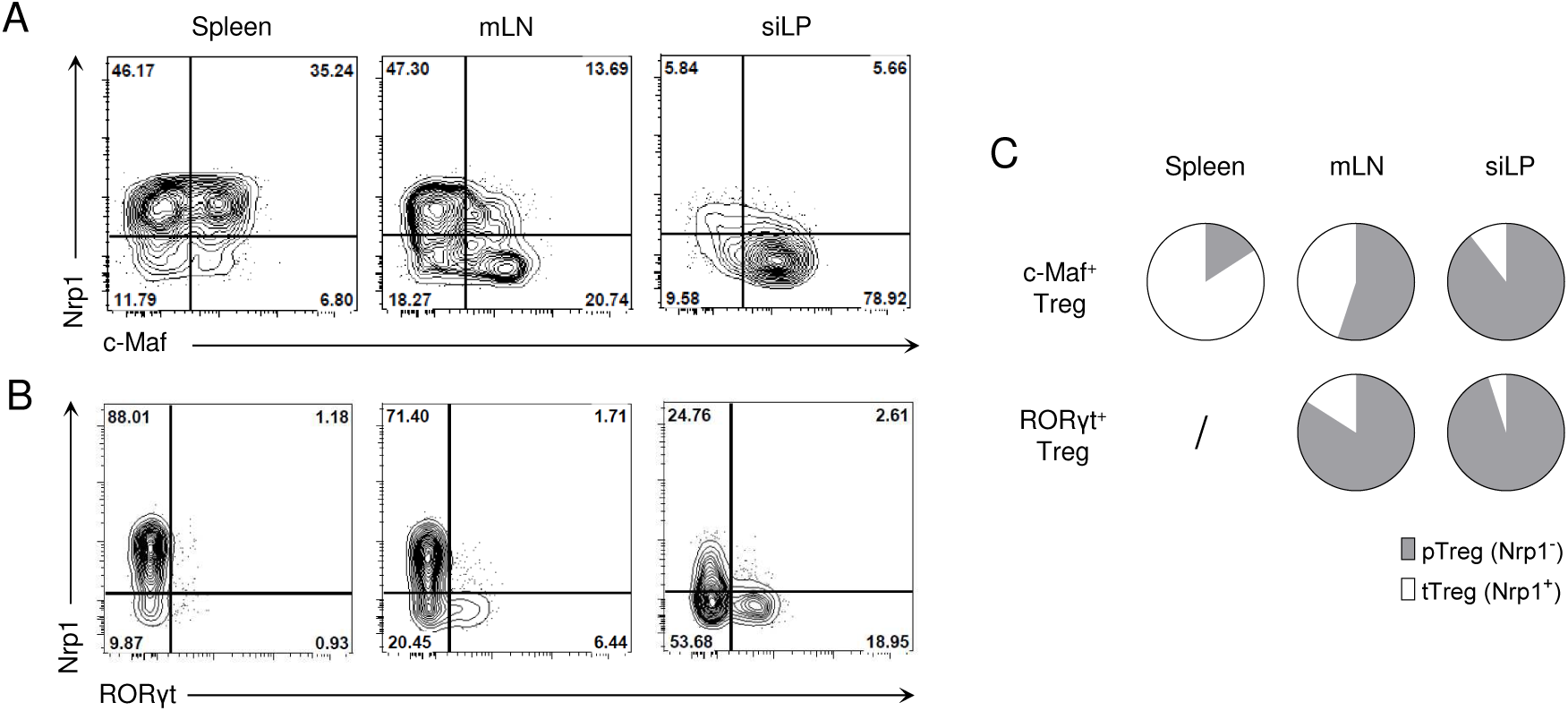
c-Maf-expressing Tregs can be of thymic or peripheral origin. (A, B) Representative flow cytometry expression profiles of Nrp1 versus c-Maf (A) or RORγt (B) of Treg cells in the indicated organs of naïve C57BL/6 mice gate CD4^+^ Foxp3^+^ cells). (C) Pie charts show the relative frequencies of thymic (tTreg) and peripheral (pTreg) Treg cells mong c-Maf^+^ and RORγt^+^ Treg subsets in the spleen, mLN and siLP. Results are representative of at least three ndependent experiments.

Treg-specific ablation of c-Maf resulted in increased Treg proportions in the intestine and lymphoid organs (Figure 1F, H and Figure S2A). Despite the wide distribution of c-Maf-expressing Tregs in distinct organs, Treg-conditional ablation of c-Maf did not result in systemic autoimmune disease, nor did it disturb conventional and regulatory T cell homeostasis in lymphoid organs. c-Maf-deficient Tregs also retained their *in vitro* suppressive capacity (Figure S2B and data not shown). In agreement with previous reports (Wheaton et al., 2017; Xu et al., 2018), c-Maf^Treg KO^ mice spontaneously developed intestinal inflammation and showed a near complete loss of RORγt expression in Tregs (Figure 1G, I and Figure S3). They nevertheless expressed normal percentages of intestinal GATA3^+^ Tregs (Figure 1G, J). Altogether, these data indicate that c-Maf is required for the differentiation of RORγt^+^ Tregs and that, contrary to RORγt, c-Maf expression is also found in a large proportion of effector thymic Tregs, located in distinct organs.

### Complex microbiota, STAT3 and TGF-β signals control RORγt but are dispensable for c-Maf expression in Tregs

The presence of c-Maf^+^ RORγt^-^ Tregs in different organs and their distinct origins prompted us to further analyze the specific environmental signals responsible for the induction of c-Maf and RORγt expression by Tregs. IL-6/STAT3 and TGF-β signaling has been shown to drive RORγt^+^ and c-Maf expression in a variety of T cells (Ohnmacht et al., 2015; Veldhoen et al., 2006). Mice genetically invalidated for IL-6 (IL-6^-/-^), STAT3 (Stat3^flox/flox^-CD4^CRE^) or TGF-β (Tgfbr2^flox/flox^-Foxp3^CRE^) signaling showed normal to even slightly increased frequencies of c-Maf^+^ Tregs, although RORγt expression was considerably decreased in all the aforementioned conditions (Figure 3; see also Figure S4 for representative FACS plots). Thus, despite being necessary for the expression of RORγt by Tregs, c-Maf is not sufficient, and most likely cooperates with other signaling pathways to induce RORγt expression in Tregs.

**Figure 3.**
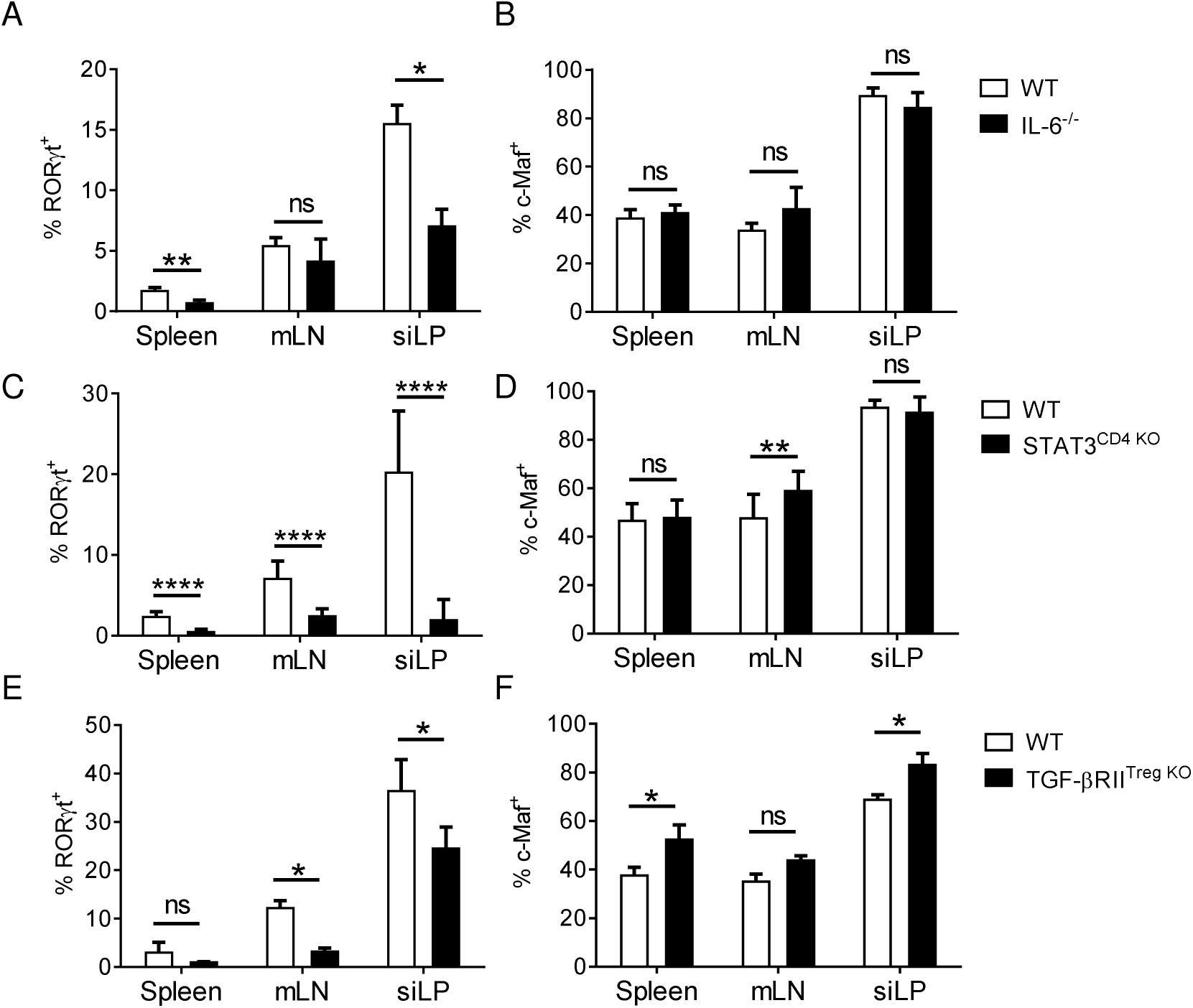
IL-6/STAT3 and TGF-β signaling promote RORγt expression in Tregs independently of c-Maf. (A-F) Histograms show the frequency of RORγt^+^ (A, C, E) or c-Maf^+^ (B, D, F) cells among Treg cells (gate CD4^+^ Foxp3^+^) in indicated organs of WT and IL-6^-/-^(A, B), STAT3^CD4 KO^ (C, D) and TGF-βRII^Treg KO^ (E, F) mice. Results are representative of at least three independent experiments; histograms represent the mean ± SD of at least four individual mice. Difference between groups is determined by an unpaired t test or a Mann–Whitney test for two-tailed data (A, B, C siLP, D mLN, E, F). See figure S4 for representative dot plots. *p < 0.05; **p < 0.01; ****p < 0.0001.

RORγt^+^ Tregs are absolutely dependent on microbiota for their development (Ohnmacht et al., 2015; Sefik et al., 2015). Analysis of germ-free and antibiotics-treated mice revealed that, in contrast to RORγt expression, which was lost in both cases, c-Maf expression by Tregs was only marginally affected in microbiota-deficient mice (Figure 4A-D).

**Figure 4.**
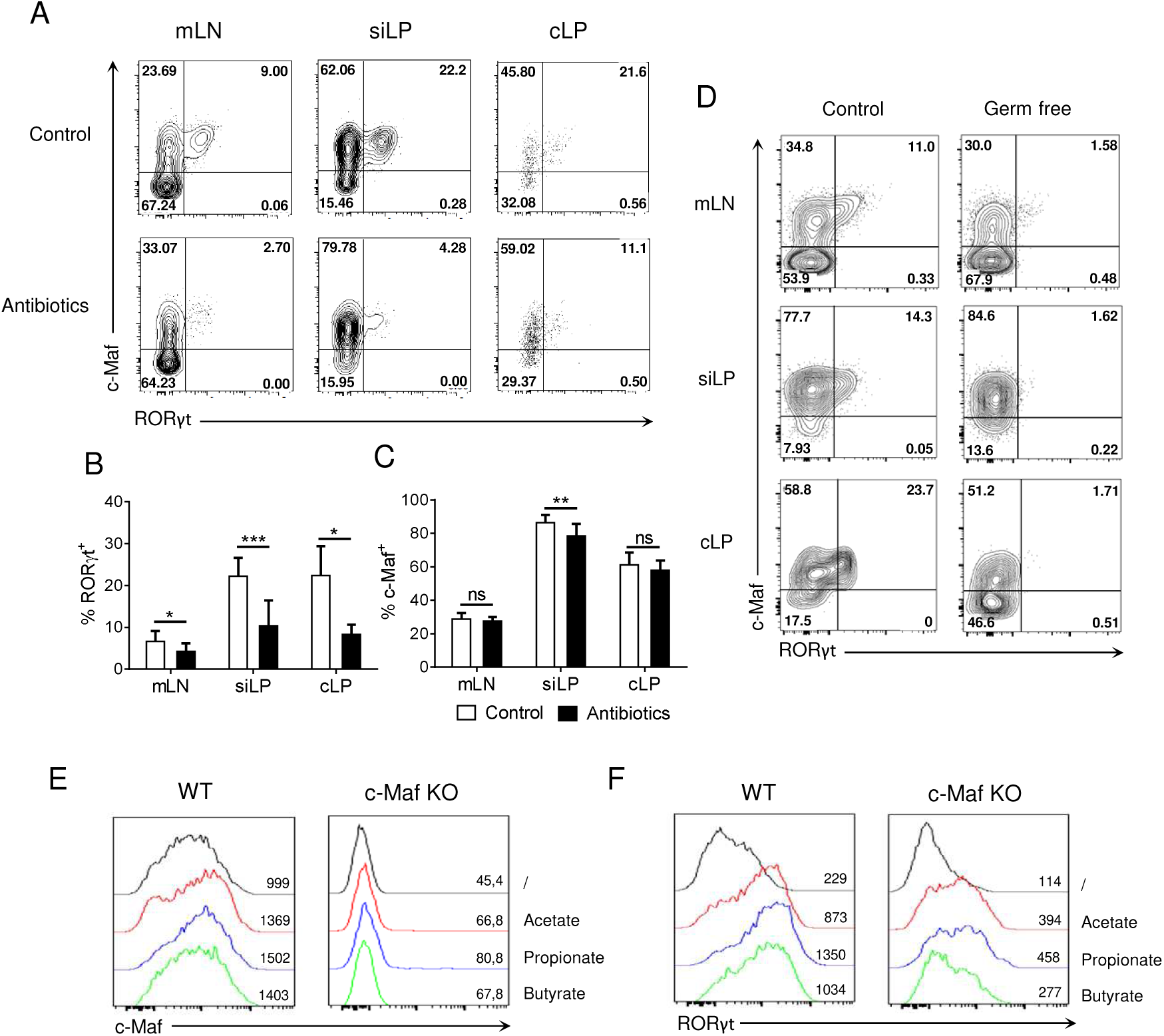
Microbial signals induce RORγt expression in Tregs but marginally affect c-Maf expression. (A-C) WT mice were treated with wide-spectrum antibiotics or a control solution for 4 weeks. Representative flow cytometry xpression profiles of c-Maf versus RORγt in Treg cells (A) and histograms showing the frequency of RORγt^+^ (B) and c-Maf^+^ (C) Tregs in the indicated organs (gate CD4^+^ Foxp3^+^). (D) Representative flow cytometry expression profiles of c-Maf versus RORγt by Treg cells in the indicated organs of germ-free and control mice. (E-F) Naïve WT or c-Maf-deficient CD4 T cells were activated *in vitro* for 72h in presence of TGF-β and small chain fatty acids. Histograms show the xpression profiles of c-Maf (F) and RORγt (G) among Treg cells. Results are representative of at least three ndependent experiments; histograms in (B, C) represent the mean ± SD of at least four individual mice. Difference etween groups is determined by an unpaired t test (mLN, siLP) or a Mann–Whitney test for two-tailed data (cLP). *p < .05; **p < 0.01; ***p < 0.001

Short chain fatty acids (SCFA) are gut microbiota-derived bacterial fermentation products, which include acetate, propionate, and butyrate, that regulate the size and function of the colonic Treg pool (Arpaia et al., 2013). Treg cells were induced *in vitro* with TGF-β in the absence or presence of acetate, propionate, or butyrate. In this experimental setting, SCFA did not affect or slightly decreased the differentiation of Foxp3^+^ Treg cells (Figure S5). Addition of SCFA to the culture medium induced a 4 to 5-fold upregulation of RORγt expression, while minimally affecting c-Maf expression (Figure 4E, F, left panels). In absence of c-Maf, intermediate levels of RORγt were induced in Tregs treated with SCFA (Figure 4F, right panel). Overall, these observations suggest that microbiota-derived products regulate RORγt expression in Tregs through both c-Maf-dependent and independent pathways.

### STAT3 and c-Maf control RORγt expression in iTregs through partly overlapping pathways

In presence of TGF-β and IL-2, about 75% of *in vitro* activated naïve CD4 T cells differentiated into Tregs, as assessed by their Foxp3 expression. Addition of IL-6, a pro-Th17 cytokine, to the TGF-β/IL-2 cytokine cocktail decreased the frequency of induced Tregs but led to the differentiation of a population of double positive c-Maf^+^ RORγt^+^ induced Tregs (iTreg17 cells; Figure 5A, lower panels and B, C). In agreement with *in vivo* observations (Figure 1G, I), ablation of c-Maf expression led to a significant reduction of *in vitro* generated RORγt^+^ Tregs (Figure 5D, E, H). In the absence of STAT3, the expression of c-Maf and RORγt in iTreg17 was heavily decreased (Figure 5F-I). Despite showing residual c-Maf expression, STAT3 KO iTregs displayed a more severe down-regulation of RORγt than their c-Maf KO counterparts (90% versus 50 %; Figure 5J), suggesting a prominent role of STAT3 in RORγt expression in this context. Analysis of STAT3 and c-Maf genome mapping from public ChIPseq databases (Ciofani et al., 2012) revealed that both transcription factors bind to the *Rorc* locus, albeit at distinct preferential sites (Figure 5K).

**Figure 5.**
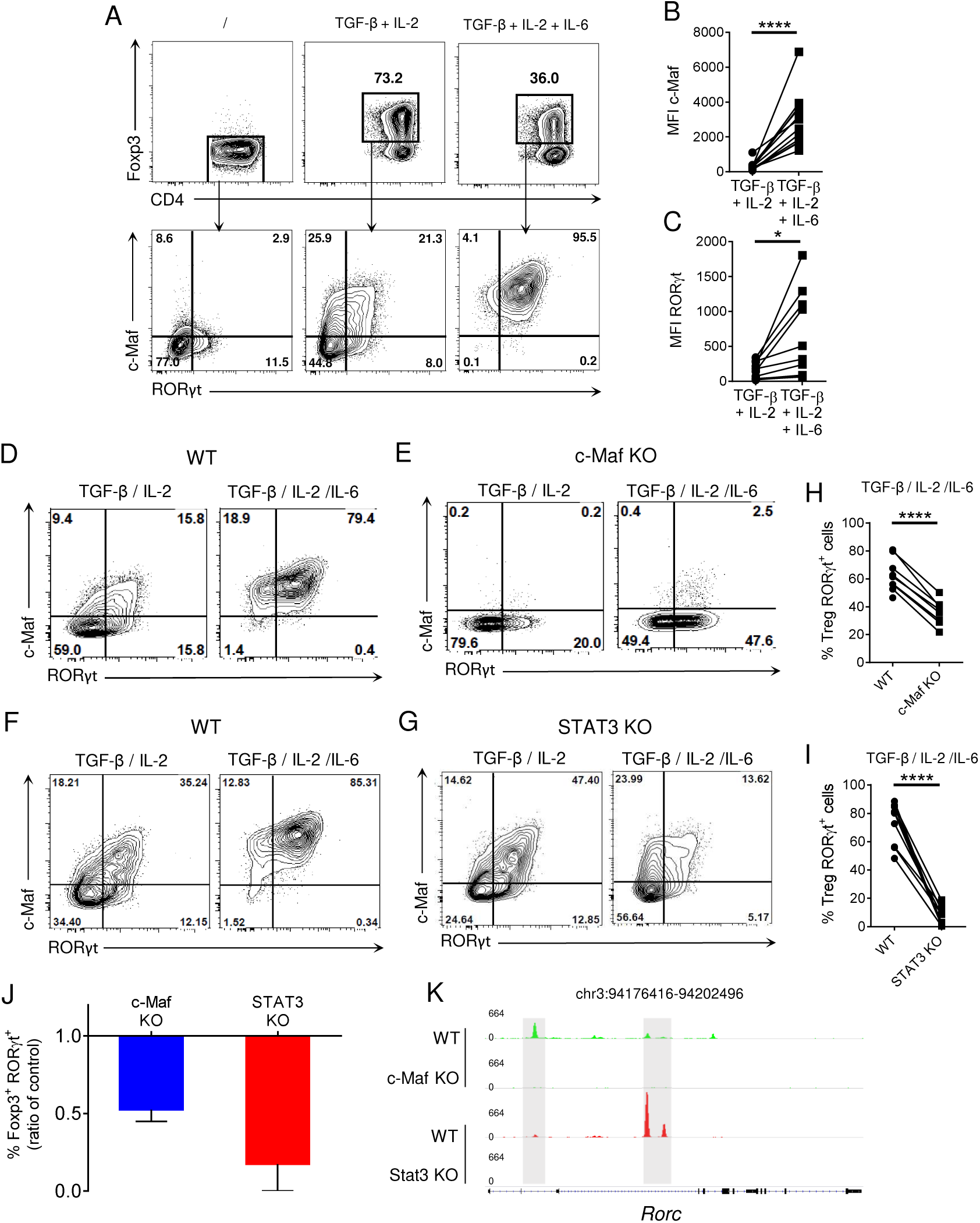
c-Maf and STAT3 are required for optimal RORγt expression in *in vitro* polarized Tregs. WT, c-Maf or STAT3-deficient naïve CD4 T cells were activated *in vitro* in presence of polarizing cytokines for 72h. Protein expression was then assessed by flow cytometry. (A) Representative flow cytometry expression profiles of Foxp3 versus CD4 and -Maf versus RORγt in the indicated gate by WT Treg cells polarized *in vitro* in the indicated conditions. (B, C) Histograms show the MFI of c-Maf (B) or RORγt (C) in WT Treg cells *in vitro*. (D-G) Representative flow cytometry expression profiles of c-Maf versus RORγt by WT (D, F), c-Maf-deficient (E) and STAT3-deficient (G) Treg cells polarized in presence of TGF-β and IL-2 or TGF-β, IL-2 and IL-6. (H, I) Histograms show the frequency of RORγt^+^ cells among WT, c-Maf (H) and STAT3-deficient (I) Tregs polarized in presence of TGF-β, IL-2 and IL-6. (J) Proportions of RORγt-xpressing cells among CD4^+^ Foxp3^+^ Tregs is expressed as a ratio of control. Values from c-Maf or STAT3-KO Tregs were divided by the value from WT Tregs in each experimental data set. (K) Profiles generated from c-Maf and STAT3 ChIP-seq in WT, c-Maf- and STAT3-KO *in vitro* Th17 cells. Representative IGV tracks showing c-Maf (green), Stat3 (red) inding sites highlighted in grey at the Rorc locus among the indicated cell population. Gene location is indicated at the op of the panel. Y axis indicates the normalized read coverage for each track. Results are representative of at least three independent experiments. Symbols in histograms represent individual mice. Difference between groups is determined by a paired t test. *p < 0.05; ****p < 0.0001.

To determine whether the role of STAT3 in promoting RORγt induction solely relies on c-Maf, we restored c-Maf expression in STAT3-deficient iTreg17 cells. WT and c-Maf KO CD4 T cells were infected with a control-GFP or a c-Maf-GFP encoding retrovirus. Although the c-Maf encoded retrovirus did restore RORγt expression in c-Maf-deficient Tregs and induced optimal levels of c-Maf in STAT3 KO Tregs, it was unable to restore RORγt expression in the latter cells (Figure 6). Collectively, these data indicate that in Tregs, an additional STAT3-driven but cMaf-independent pathway is required to promote optimal RORγt.

**Figure 6.**
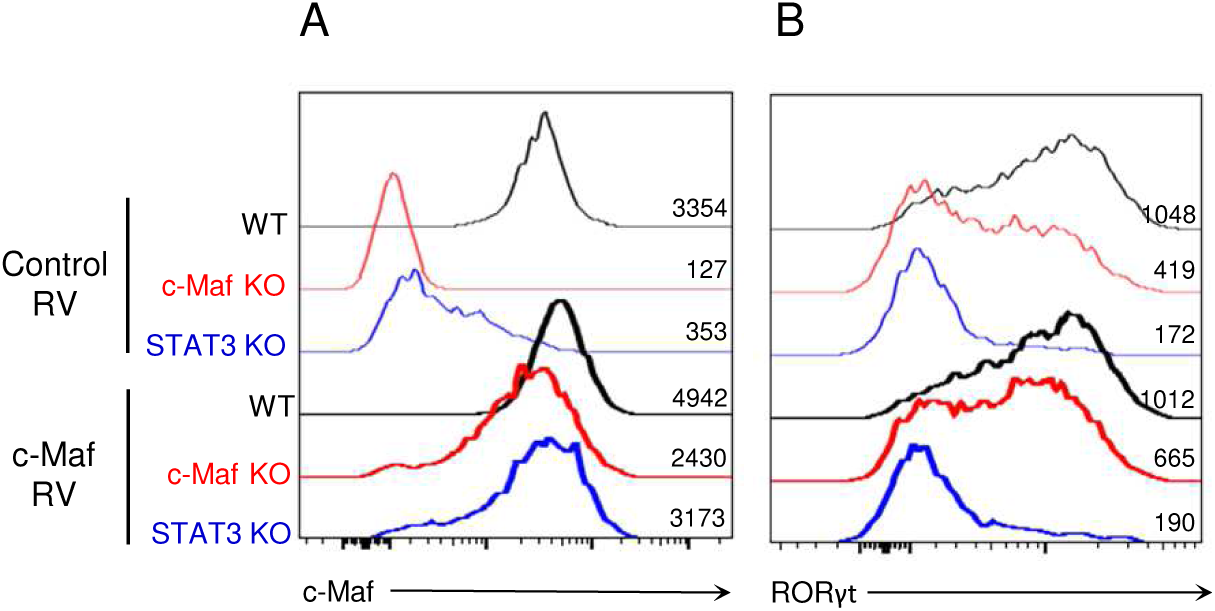
c-Maf does not rescue RORγt expression in Tregs in the absence of STAT3. WT, c-Maf or STAT3-deficient aïve CD4 T cells were activated *in vitro* in presence of TGF-β, IL-2 and IL-6 and infected with a control or c-Maf-xpressing retrovirus. c-Maf (A) and RORγt (B) expression profiles and MFI as assessed by flow cytometry after 72h gate CD4^+^ Foxp3^+^). Results are representative of three independent experiments

### A pro-Th1 inflammatory environment dampens RORγt expression in Tregs

While IL-6 promoted RORγt expression in wild type iTregs, it surprisingly led to a marked reduction of this transcription factor in STAT3-deficient iTregs (Figure 5F-I), thus revealing the presence of an inhibitory pathway regulating RORγt expression. Studies by Costa-Pereira et al have shown that in STAT3 KO fibroblasts, IL-6 signals through STAT1 and has IFNγ-like effects (Costa-Pereira et al., 2002). We observed that in contrast to their wild type counterparts, STAT3 KO CD4^+^ T cells cultured in the presence of IL-6 expressed higher levels of phospho-STAT1 (Figure S6). This prompted us to investigate the consequences of a STAT1/Th1 inflammatory pathway on the differentiation of RORγt^+^ Tregs.

Infection with *Toxoplasma gondii* results in a highly Th1-polarized microenvironment leading to altered Treg cell homeostasis (Oldenhove, 2009). In particular, the strong Th1 environment triggered by *T. gondii* infection has been shown to induce T-bet and IFN-γ expression in Treg cells (Hall et al., 2012; Oldenhove et al., 2009). Analysis of Tregs from the lamina propria of *T. gondii* infected C57BL/6 mice confirmed the emergence of a T-bet^+^ Treg subset (Figure 7A and B, upper panels). Interestingly, *T. gondii* infection resulted in a reduced frequency of tissue associated RORγt^+^ Tregs, despite having minimal effect on Maf expression (Figure 7A and B, middle and lower panels). Th17 cell differentiation was also suppressed during *T. gondii* infection (Figure S7), consistent with the observation that Th17 cell differentiation and Th1 cell differentiation are mutually suppressive (Harrington et al., 2005; Park et al., 2005). This reciprocal exclusion was also reflected among Tregs, as T-bet and RORγt were expressed in distinct Treg cell subsets in the infected mice (Figure 7C). We next wished to evaluate the effect of Th1-promoting signals on RORγt expression by Tregs, Addition of IFN-γ to the iTreg17 differentiation media led to the selective accumulation of the phosphorylated form of STAT1, without affecting neither phospho-STAT3 accumulation nor c-Maf expression by iTregs (Figure S8). Of note, IFN-γ led to the accumulation of Tbet-expressing Tregs, with a concomitant reduction in the number of RORγt^+^ Tregs (Figure 7D, upper panels). STAT1 KO Tregs were insensitive to the IFN-γ-driven inhibition of RORγt expression (Figure 7D, lower panels). Altogether these results strongly suggest that the IFN-γ/STAT1 signaling pathway negatively regulates RORγt expression even in the presence of an active IL-6/STAT3-driven pathway.

**Figure 7.**
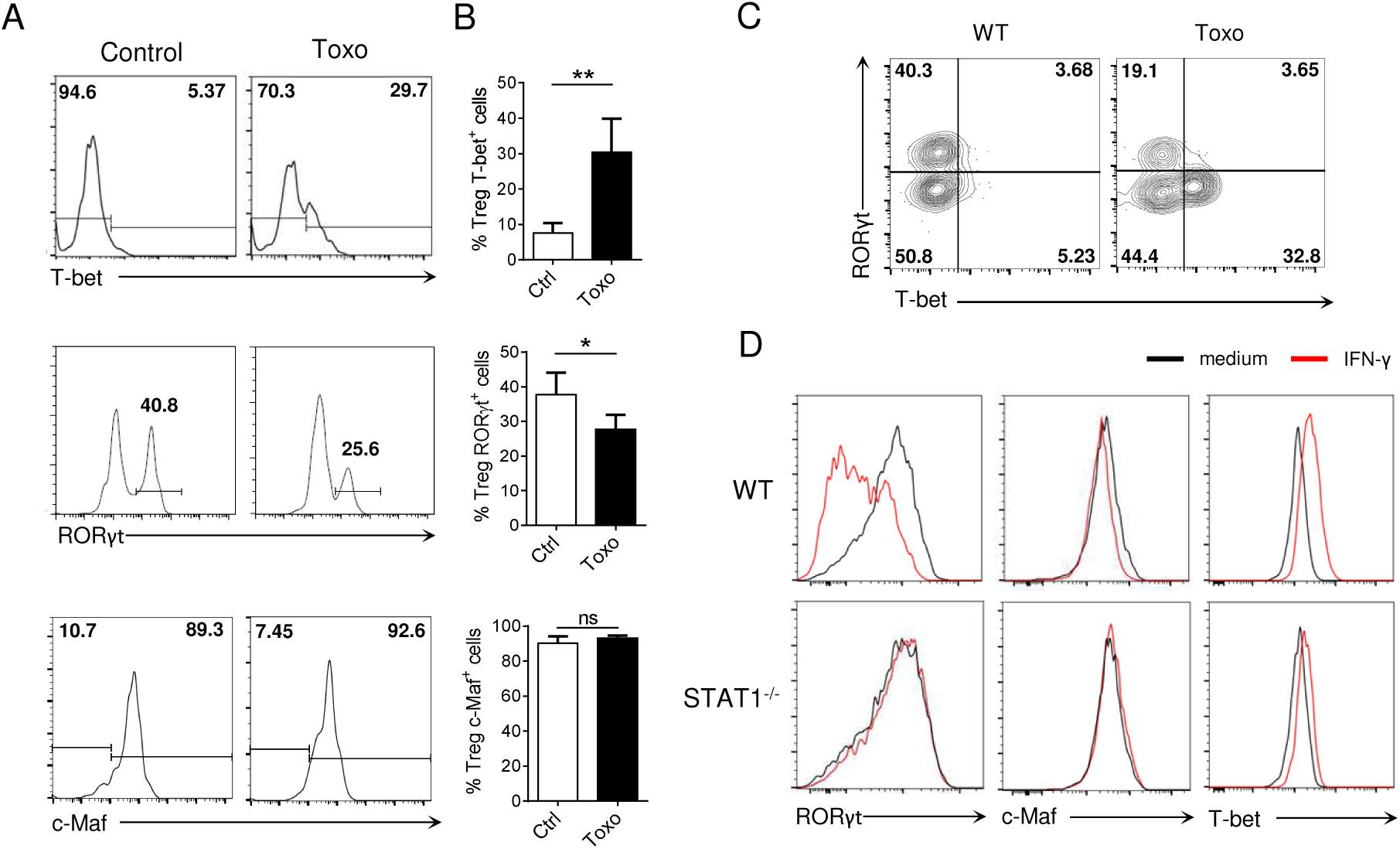
Inflammatory Th1 responses opposes RORγt expression in Tregs. (A) Histograms show RORγt, c-Maf nd T-bet expression among Tregs in the siLP of WT mice 8 days after infection with *Toxoplasma gondii* or control mice gate CD4^+^ Foxp3^+^). (B) Frequency of RORγt^+^, c-Maf^+^ and T-bet^+^ cells among Treg cells in the siLP of WT mice 8 days after infection with *Toxoplasma gondii* or control mice. (C) Representative flow cytometry expression profiles of RORγt ersus T-bet among Treg cells isolated from mLN of WT mice infected or not with *Toxoplasma gondii* (gate CD4^+^ Foxp3^+^). (D) Histograms show RORγt, c-Maf and T-bet expression in WT and STAT1-deficient *in vitro* Treg17 cells stimulated with or without IFN-γ (100 ng/mL) for 72h. Results are representative of at least two (A-C) or three (D) independent experiments; histograms represent the mean ± SD of five individual mice. Difference between groups is determined by a Mann–Whitney test for two-tailed data. *p < 0.05

## Discussion

RORγt^+^ Tregs form a distinct population of regulatory T cells that is crucial to maintain gastrointestinal homeostasis and prevent colitis (Ohnmacht et al., 2015; Sefik et al., 2015; Yang et al., 2015). Whereas RORγt^+^ Treg function has been amply documented, the factors driving RORγt^+^ Treg differentiation remain ill-defined. We show herein that multiple signaling pathways cooperate to exert both positive and negative control over RORγt^+^ expression by Tregs.

We confirmed and extended previous data showing that transcription factor c-Maf plays a major role for the acquisition of RORγt expression by Tregs (Neumann et al., 2019; Wheaton et al., 2017; Xu et al., 2018). However, contrary to RORγt, which is confined to a subset of intestinal Tregs, c-Maf expression is found in a wider proportion of Tregs, located in distinct organs. This observation suggests that c-Maf cooperates with other pathways to induce optimal RORγt expression by these regulatory cells.

We showed that microbial signals, as well as IL-6-, STAT3-, and TGF-β-signaling pathways promote RORγt^+^ Treg cell differentiation in a non-redundant manner. Constrastingly, blocking any one of these pathways only had minimal effect on c-Maf expression *in vivo*, suggesting that several redundant pathways cooperate to induce c-Maf expression in Tregs. Neumann et al reported that, in addition to the loss of RORγt expression in Tregs, which we also observed, germ-free mice exhibited a near complete loss of c-Maf expression in intestinal Tregs. While we could not reproduce this observation, we observed that Tregs found in germ-free mice expressed slightly reduced levels of c-Maf. Reminding that the RORγt^+^ population expresses the highest levels of c-Maf among intestinal Tregs, we therefore speculate that RORγt expression in Tregs requires a high c-Maf threshold and that even a minimal decrease in c-Maf expression could hinder RORγt expression in microbiota-driven intestinal Tregs. Short chain fatty acids (SCFA) produced by gut commensal microbes induce functional colonic Tregs and protect against T cell-dependent experimental colitis (Arpaia et al., 2013; Furusawa et al., 2013). Depending on the cytokine environment and immunological context, butyrate, acetate, and propionate, the most available SCFA in the gut, can also support IL-10 expression in Th1 and Th17 effector cells, thereby inhibiting colitis caused by pathogenic T cells (Park et al., 2015; Sun et al., 2018). We show herein that SCFA induce RORγt expression in *in vitro* differentiated Tregs in a c-Maf-dependent and independent manner. Further studies could help clarify whether SCFA regulate RORγt expression through GPR41 or GPR43-dependent signaling, or through HDAC inhibitor activity and subsequent enhancement of mTOR–S6K activity, as previously shown in Th17 cells (Park et al., 2015). The proportion of RORγt^+^ Tregs was severely reduced in mice deficient for STAT3 or TGF-β signaling. Rather surprisingly, while STAT3 and TGF-β signaling drive c-Maf expression in Tfh and Th17 cells (Ciofani et al., 2012; Mari et al., 2013; Rutz et al., 2011), we found that mice deficient for STAT3 or TGF-β in the T cell compartment harbored normal to even slightly increased numbers of c-Maf^+^ Tregs, indicating that, while STAT3 and TFG-β seem dispensable for the induction of c-Maf expression, they are both essential to achieve optimal differentiation of RORγt^+^ Tregs in the intestine.

Besides c-Maf, many genes involved in Th17 differentiation, such as Irf4, Batf, Rora, Ahr, Sox5t and HIF-1α, are expressed in response to STAT3 activation (Brüstle et al., 2007; Dang et al., 2011; Durant et al., 2010; Schraml et al., 2009; Tanaka et al., 2014b; Yang et al., 2008). Sox5t is a T cell isoform of Sox5 which induces RORγt expression in Th17 cells via physical interaction with c-Maf (Tanaka et al., 2014b). As enforced expression of c-Maf was not sufficient to induce Foxp3^+^ RORγt^+^ T cell differentiation in absence of STAT3, we speculate that Sox5, or other molecules downstream of STAT3, act together with c-Maf to achieve RORγt expression in Tregs. Interestingly, ChIP-seq data revealed that STAT3 and c-Maf bind with different intensities to distinct sites of the *Rorc* locus. The inability of c-Maf to compensate STAT3-deficiency in RORγt^+^ Treg differentiation could thus also be explained by a direct effect of STAT3 on RORγt expression. This is in agreement with previous data showing that STAT3 binds to intron 1 of *Rorc* gene and induces chromatin remodeling of the locus (Durant et al., 2010). Overall, it would seem that STAT3 controls RORγt^+^ Treg cell fate both through direct activation of the *Rorc* locus and by regulating the expression of a set of genes, including c-Maf, that is essential for RORγt^+^ Treg differentiation (Figure S9).

In T cells, IL-6 predominantly signals via STAT3 and to a lesser extent via STAT1. It has been proposed that the accessibility of different STATs within the cell influences the outcome of cytokine signaling (Regis et al., 2008), as illustrated by the observation that IL-6 acquired the ability to induce the expression of STAT1-dependent genes in STAT3-deficient cells (Costa-Pereira et al., 2002; Schiavone et al., 2011). However, Hirahara et al showed an asymmetric action of STAT3 and STAT1 at the genomic level where much of STAT1 chromatin binding was STAT3-dependent. This challenged the classical view that, in the absence of its major STAT module, a cytokine would acquire an alternative STAT-signaling profile (Hirahara et al., 2015). Yet, in the absence of STAT3, and despite a global reduction in STAT1 chromatin binding, some preferential STAT1 binding sites were conserved in a group of IL-6 downregulated genes. With this in mind we hypothesized a negative influence of STAT1 on RORγt expression by Tregs. Indeed, STAT3-deficient T cells showed increased STAT1 phosphorylation in response to IL-6, compatible with a switch from STAT3 to STAT1 signaling in these cells. We further showed that IFN-γ-driven activation of STAT1 opposed RORγt expression in Tregs, without affecting Foxp3 and c-Maf expression or STAT3 phosphorylation. Naïve STAT1 KO mice did not show altered proportions of intestinal RORγt^+^ Tregs (data not shown), which could be explained by the lack of inflammatory Th1 components at steady state. Indeed, in wild type mice infected with the Th1-prototypic *Toxoplasma gondii* intestinal parasite, RORγt expression was decreased in intestinal Tregs, confirming the antagonistic role of inflammatory Th1 responses on RORγt expression in Tregs *in vivo*.

Different integrative pathways have been proposed to explain the functional outcome of multiple STAT signaling in distinct T cell subsets (Lin and Leonard, 2019). Meyer zu Horste et al recently reported that death receptor Fas promotes Th17 cell differentiation and inhibits Th1 cell development by preventing STAT1 activation. In this model, Fas regulated the STAT1 versus STAT3 balance by binding and sequestering STAT1 (Meyer zu Horste et al., 2018). Although not formerly excluded, sequestration of STAT3 is unlikely in RORγt^+^ Tregs as addition of IFN-γ to the iTreg17 culture media did not affect the phosphorylation status of STAT3. Our data rather suggest that Treg cell fate results from the balance of STAT1 and STAT3 driven signals. Gene expression could also conceivably be fine-tuned by the formation of STAT1/STAT3 heterodimers, as proposed for the IL-21 signaling (Wan et al., 2015). Of interest, patients with STAT3 mutations or with gain-of-function STAT1 mutations show similar susceptibility to fungal infections (Casanova et al., 2012; O’Shea et al., 2013). In the latter group, overactive STAT1 appears to limit STAT3-driven antifungal responses (Casanova et al., 2012).

Further work is required to decipher whether STAT1 interacts with STAT3 or exerts an independent negative role on RORγt^+^ Treg cell fate. As T-bet is induced in Tregs that develop during *Toxoplasma* infection or in response to IFN-γ, we can also envision that STAT1 signaling inhibits RORγt expression through T-bet blocking of Runx1-mediated transactivation of *Rorc*, as previously reported for the Th1/Th17 lineage specification (Lazarevic et al., 2011). Regardless of the molecular mechanism, the observation that IFN-γ/STAT1 signaling pathway negatively regulates RORγt, even in the presence of an active IL-6/STAT3 pathway, suggests a dominant negative effect of STAT1 over STAT3 in these experimental conditions.

The antagonism between STAT1 and STAT3 seems to be cell type-specific or specific to a certain gene locus, as a cooperation between STAT1 and STAT3 downstream of IL-6 has been described for optimal Bcl6 induction and Tfh differentiation in response to viral infections (Choi et al., 2013).

Collectively, our data reveal that, beyond the previously established c-Maf/RORγt interplay, multiple signaling pathways cooperate to exert a tight control over RORγt expression in Tregs.

## Material and Methods

### Mice

C57BL/6 mice were purchased from Envigo (Horst, The Netherlands). c-Maf-flox mice (C. Birchmeier, Max Delbrück Center for Molecular Medicine, Berlin, Germany) were crossed with CD4-CRE mice (G. Van Loo, Ghent University, Ghent, Belgium) or FOXP3-CRE-YFP mice which were developed by A. Rudensky (Rubtsov et al., 2008) and kindly provided by A. Liston (KU Leuven, Leuven, Belgium). IL-6^-/-^ mice were obtained from The Jackson Laboratory (Bar Harbor, ME, USA). STAT3-flox mice were kindly provided by S. Akira (Osaka University, Osaka, Japan); STAT1^-/-^ mice by D.E. Levy (New York University School of Medicine, NYC, USA). Germ-free mice were obtained from the Ghent Germfree and Gnotobiotic mouse facility (Ghent University, Ghent, Belgium) and were compared to SPF control mice. c-Maf-flox, CD4-CRE, FOXP3-CRE-YFP, IL-6^-/-^, STAT3-flox, STAT1^-/-^ and germ-free mice were bred on a C57BL/6 background. Tgfbr2-flox mice (Chytil et al., 2002) on a NOD background crossed with Foxp3-Cre mice (JAX 008694) were kindly provided by Q. Tang (University of California San Francisco, SF, USA) and were housed and bred at the UCSF Animal Barrier Facility.

All mice were used between 6 and 12 weeks of age. The experiments were carried out in compliance with the relevant laws and institutional guidelines and were approved by the Université Libre de Bruxelles Institutional Animal Care and Use Committee (protocol number CEBEA-4).

### Antibodies, intracellular staining and flow cytometry

The following monoclonal antibodies were purchased from eBioscience: CD278 (ICOS)-biotin, CD304 (Nrp1)-PerCP eF710, c-Maf-EF660, Foxp3-FITC, RORγt-PE, T-bet-PE; or from BD Biosciences: CD25-BB515, CD44-PECy7, CD4-A700, CD4-PB, CD62L-A700, GATA3-PE, RORγt-PECF594, STAT1 (pY701)-A488, STAT3 (pY705)-A647, streptavidin-PECy7.

Live/dead fixable near-IR stain (ThermoFisher) was used to exclude dead cells. For transcription factor staining, cells were stained for surface markers, followed by fixation and permeabilization before nuclear factor staining according to the manufacturer’s protocol (FOXP3 staining buffer set from eBioscience). For phosphorylation staining, cells were fixed with formaldehyde and permeabilized with methanol before staining. Flow cytometric analysis was performed on a Canto II (BD Biosciences) or CytoFLEX (Beckman Coulter) and analyzed using FlowJo software (Tree Star).

### Isolation of lymphocytes

After removal of Peyer’s patches and mesenteric fat, intestinal tissues were washed in HBSS 3% FCS and PBS, cut in small sections and incubated in HBSS 3% FCS containing 2,5mM EDTA and 72,5 µg/mL DTT for 30 min at 37°C with agitation to remove epithelial cells, and then minced and dissociated in RPMI containing liberase (20 µg/ml, Roche) and DNase (400 µg/ml, Roche) at 37 °C for 30 min (small intestine) or 45 min (colon). Leukocytes were collected after a 30% Percoll gradient (GE Healthcare). Lymph nodes, thymus and spleens were mechanically disrupted in culture medium.

After anesthesia, mice were perfused with PBS. Liver and lung samples were digested with collagenase (200U, Worthington) and DNase I (40µg/mL, Roche) at 37°C and mechanically disrupted. Leukocytes were collected at the interphase of a 40%/70% Percoll gradient.

### Antibiotics treatment

Wide spectrum antibiotics (ampicillin 1g/L and neomycin 1g/L, Sigma-Aldrich; vancomycin 0,5g/L and metronidazole 1 g/L, Duchefa) were added to the sweetened drinking water of mice treated with antibiotics for three to four weeks. Control mice were given sweetened drinking water in parallel.

### Toxoplasma infection

ME-49 type II *T. gondii* was kindly provided by Dr De Craeye (Institut Scientifique de Santé Publique, Belgium) and was used for the production of tissue cysts in C57BL/6 mice, which were inoculated 1–3 months previously with three cysts by gavage. Animals were killed, and the brains were removed. Tissue cysts were counted and mice were infected by intragastric gavage with 10 cysts. Mice were sacrificed at day 8 after infection.

### T cell culture

Naive CD4^+^ T cells were purified from spleen of mice with indicated genotypes. CD4^+^ T cells were positively selected from organ cell suspensions by magnetic-activated cell sorting using CD4 beads (MACS, Miltenyi) according to the product protocol, and then isolated as CD4^+^ CD44^lo^CD62L^hi^CD25^−^ or CD4^+^ CD44^lo^ CD62L^hi^ YFP^−^ by FACS. T cells were cultured at 37°C in RPMI supplemented with 5% heat-inactivated FBS (Sigma-Aldrich), 1% non-essential amino acids (Invitrogen), 1 mM sodium pyruvate (Invitrogen), 2 mM L-glutamin (Invitrogen), 500 U/mL penicillin/500 µg/ml streptomycin (Invitrogen), and 50 μM β-mercaptoethanol (Sigma-Aldrich).

To generate iTreg cells, cells were cultured 72h in 24 or 96 well plates coated with 5µg/mL anti-CD3 (BioXcell, 145-2c11) at 37°C for 72h. The culture was supplemented with anti-CD28 (1 µg/mL, BioXcell, 37.51) and TGF-β alone (3 ng/ml, eBioscience), TGF-β and IL-2 (10 ng/mL, Peprotech), or TGF-β, IL-2 and IL-6 (10 ng/mL, eBioscience) for optimal iTreg cell polarization. Acetate (C2, 10 mM), propionate (C3, 0,5 mM), butyrate (C4, 0,125 mM), all from Sigma-Aldrich, and IFN-γ (10 and 100 ng/mL, Peprotech) were also used and added to the culture for the whole duration of the experiment.

### Retroviral infection

Platinum-E retroviral packaging cells (T. Kitamura, University of Tokyo, Tokyo, Japan) were transfected with a c-Maf encoding retroviral plasmid (pMIEG-c-Maf-IRES-GFP) or a control retroviral plasmid (pMIEG-IRES-GFP) to produce retrovirus-containing supernatants. 24 hours after activation, naïve CD4 T cells polarized in presence of TGF-β, IL-2 and IL-6, as described above, were infected during a 90-minute centrifugation with 1 mL retrovirus-containing supernatant and polybrene. 48 hours later, infected cells were FACS-sorted based on GFP expression and were stimulated with anti-CD3 for 24 hours (5µg/mL, coated) before flow cytometry staining.

### Treg cell *in vitro* suppression assay

Naïve T cells with the phenotype CD4^+^ CD44^lo^ CD62L^hi^ YFP^−^ were isolated from the spleen of FOXP3-CRE-YFP mice by FACS after positive enrichment for CD4^+^ cells using MACS LS columns (Miltenyi) and labelled with carboxyfluorescein diacetate succinimidyl ester (CFSE, ThermoFisher). Treg cells with the phenotype CD4^+^ FOXP3-CRE-YFP^+^ were isolated from the mesenteric lymph nodes of Foxp3-CRE-YFP or c-Maf^Treg KO^ mice by FACS. Splenocytes from wild-type B6 mice were depleted in T cells (anti-CD90.2 beads) using MACS LS columns (Miltenyi). 4 × 10^4^ CFSE-labelled naive T cells were cultured for 72 h with APCs (1 × 10^5^) and soluble anti-CD3 (0,5 μg/mL) in the presence or absence of various numbers of Treg cells as indicated.

### RT-qPCR

RNA was extracted using the TRIzol method (Invitrogen) and reverse transcribed with Superscript II reverse transcriptase (Invitrogen) according to the manufacturer’s instructions. Quantitative real-time RT-PCR was performed using the SYBR Green Master mix kit (ThermoFisher). Primer sequences were as follow: RPL32 (F) ACATCGGTTATGGGAGCAAC; RPL32 (R) TCCAGCTCCTTGACATTGT; IL-10 (F) CCTGGGTGAGAAGCTGAAGA; IL-10 (R) GCTCCACTGCCTTGCTCTTA; IL-17A (F) ATCCCTCAAAGCTCAGCGTGTC; IL-17A (R) GGGTCTTCATTGCGGTGGAGAG; IL-22 (F) CAGCAGCCATACATCGTCAA; IL-22 (R) GCCGGACATCTGTGTTGTTA; TNF-α (F) GCCTCCCTCTCATCAGTTCTA; TNF-α (R) GCTACGACGTGGGCTACAG.

### ChIP-seq data

Publicly available ChIP-seq data (GSE40918) for c-Maf and Stat3 was downloaded and mapped to the mm9 genome using Bowtie2 with sensitive-local predefined parameters. Resulted BAM files were converted to bigwig files and visualized by IGV genome browser.

### Statistical analysis

For unpaired data, statistical difference between groups was determined by an unpaired t test when the sample size was sufficient and both groups passed the normality test, and by a Mann-Whitney test for two-tailed data otherwise. Mutant and control groups did not always have similar standard deviations and therefore an unpaired two-sided Welch’s t-test was used. For paired data, a paired t test was used. Error bars represent mean ± SD. No samples were excluded from the analysis.

## Acknowledgments

We especially thank Muriel Moser for her great support all along this work, for stimulating discussions and for careful revision of the manuscript. We also thank Caroline Abdelaziz and Véronique Dissy for animal care and for technical support. This work was supported by the European Regional Development Fund (ERDF) and the Walloon Region (Wallonia-Biomed portfolio, 411132-957270), grant from the Fonds Jean Brachet and research credit from the National Fund for Scientific Research, FNRS, Belgium. F.A. is a Research Associate at the FNRS. Y.A. is recipient of a research fellowship from the FNRS/Télévie. H.H. is supported by a Belgian FRIA fellowship.

