## Supplemental Fig 1 for "Multiple environmental signaling pathways control the differentiation of RORγt-expressing regulatory T cells"

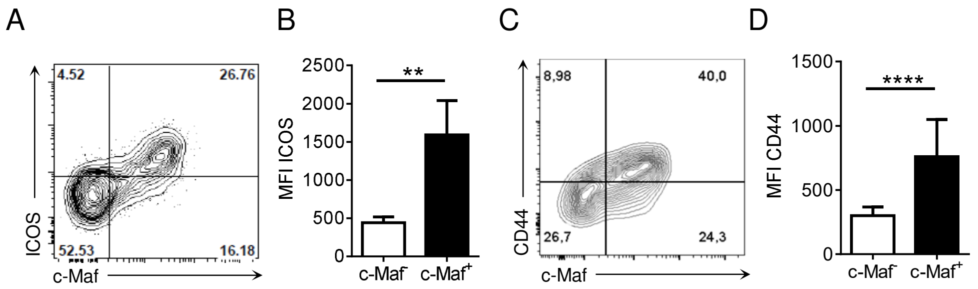


Figure S1. **c-Maf+ Tregs have an activated phenotype.** (A, C) Representative flow cytometry expression profiles of ICOS (A) or CD44 (C) versus c-Maf among Treg cells in mLN of WT mice (gate CD4+ Foxp3+). (B, D) Histograms show the MFI of ICOS (B) and CD44 (D) among c-Maf- and c-Maf+ Treg cells. Histograms represent the mean ± SD of at least five individual mice. Difference between groups is determined by a Mann–Whitney test for two-tailed data. **p < 0.01; ****p < 0.0001
