## Supplemental Fig 2 for "Multiple environmental signaling pathways control the differentiation of RORγt-expressing regulatory T cells"

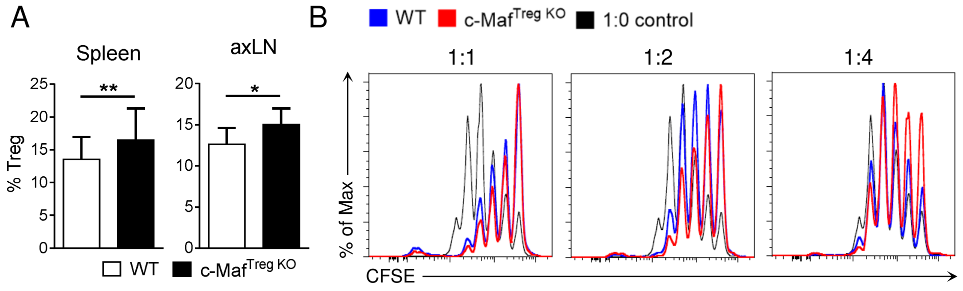


Figure S2. **c-Maf-deficient Tregs have a suppressive phenotype.** (A) Frequency of Treg cells in the spleen or axillary lymph nodes of WT and c-Maf$^{Treg\ KO}$ mice (gate CD4$^{+}$). (B) Histograms showing CFSE staining profiles of conventional CD4 T cells in a Treg suppression assay *in vitro*. Results are representative of at least three independent experiments; histograms represent the mean ± SD of five individual mice. Difference between groups is determined by a Mann–Whitney test for two-tailed data. *p < 0.05; **p < 0.01
