## Supplemental Fig 3 for "Multiple environmental signaling pathways control the differentiation of RORγt-expressing regulatory T cells"

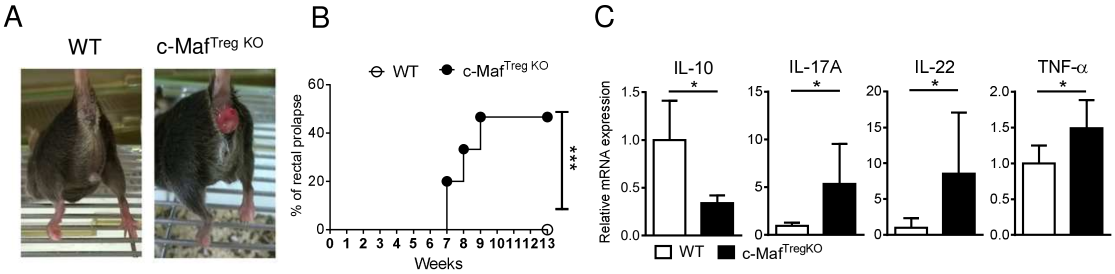


Figure S3. **c-Maf-deficient Tregs are unable to control spontaneous Th17 intestinal responses.** (A) Rectal prolapse appears in c-Maf$^{Treg\ KO}$ mice. (B) Incidence of rectal prolapse in WT and c-Maf$^{Treg\ KO}$ mice. (C) Histograms show mRNA expression of IL-10, IL-17A, IL-22 and TNF-α in total WT or c-Maf$^{Treg\ KO}$ mLN relative to RPL32. Results are representative of at least three independent experiments; histograms represent the mean ± SD of five individual mice. Difference between groups is determined by a Mantel-Cox test (B) or a Mann–Whitney test for two-tailed data. *$p < 0.05$; ***$p < 0.001$
