## Supplemental Fig 4 for "Multiple environmental signaling pathways control the differentiation of RORγt-expressing regulatory T cells"

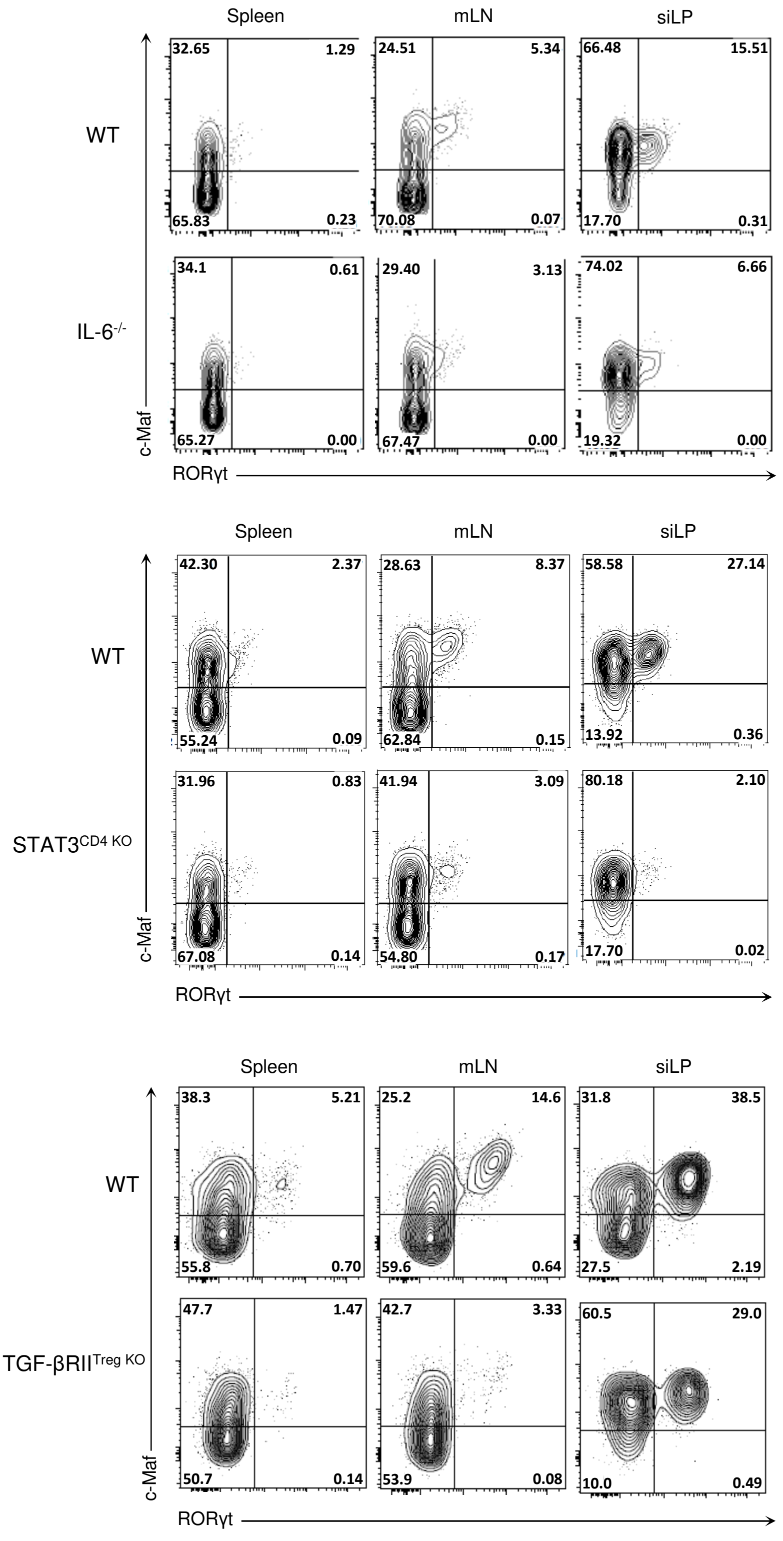


Figure S4. **IL-6/STAT3 and TGF-β signaling promote RORγt expression in Tregs independently of c-Maf**. Representative flow cytometry expression profiles of c-Maf versus RORγt in Treg cells in the indicated organs of the indicated mice strains (gate CD4+ Foxp3+). Results are representative of at least three independent experiments.
