## Supplemental Fig 5 for "Multiple environmental signaling pathways control the differentiation of RORγt-expressing regulatory T cells"

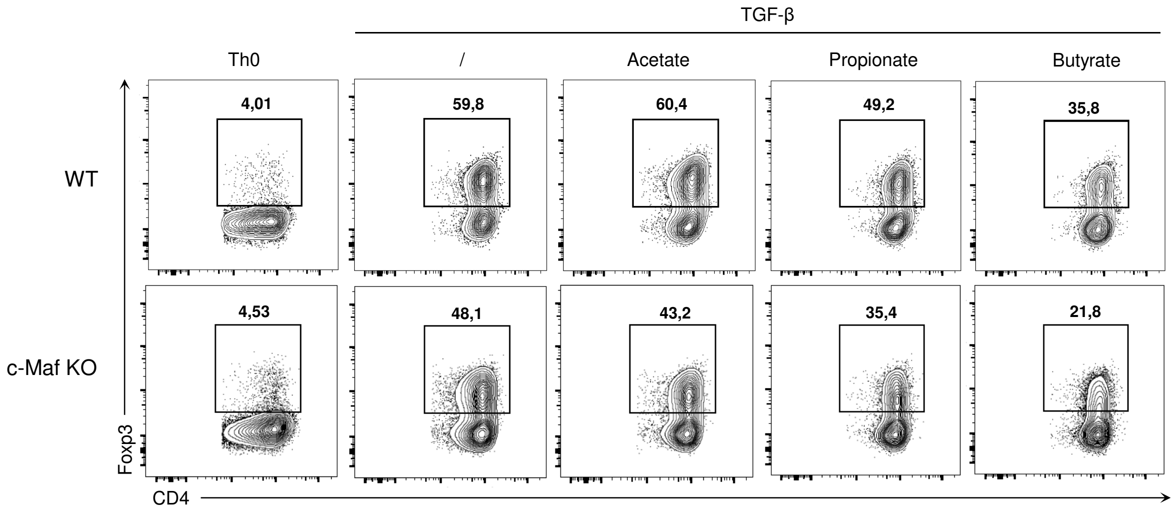


Figure S5. **In vitro Treg differentiation in presence of short-chain fatty acids.** Naïve WT or c-Maf-deficient CD4 T cells were activated *in vitro* for 72h in presence of TGF-β and small chain fatty acids. Representative flow cytometry expression profiles of Foxp3 in CD4 cells in the indicated conditions; gating strategy of Fig. 4E, F. Results are representative of at least three independent experiments.
