## Supplemental Fig 6 for "Multiple environmental signaling pathways control the differentiation of RORγt-expressing regulatory T cells"

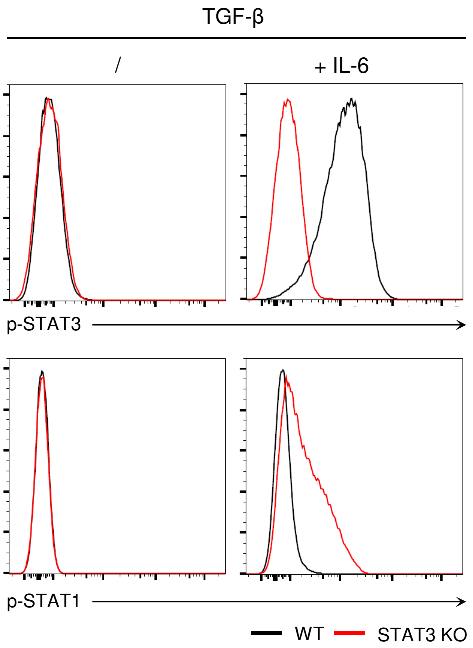


Figure S6. **In absence of STAT3 in Tregs, IL-6 signals via STAT1.** Histograms show the expression of pSTAT3 and pSTAT1 among Treg cells polarized *in vitro* in presence of TGF-β, with or without IL-6 (gate CD4+).
