## Supplemental Fig 7 for "Multiple environmental signaling pathways control the differentiation of RORγt-expressing regulatory T cells"

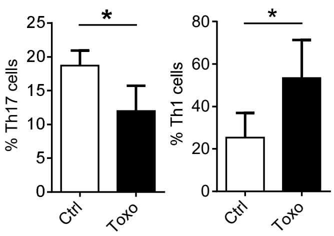


Figure S7. **Infection with *Toxoplasma gondii* is characterized by a Th1 inflammatory environment.** Histograms show the frequency of Th17 cells and Th1 cells in siLP of mice after *Toxoplasma gondii* infection. Histograms represent the mean ± SD of five individual mice. Difference between groups is determined by a Mann–Whitney test for two-tailed data. **$p < 0.01$; ****$p < 0.0001$
