## Supplemental Fig 8 for "Multiple environmental signaling pathways control the differentiation of RORγt-expressing regulatory T cells"

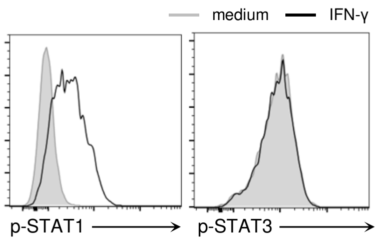


Figure S8. **IFN-γ does not impede IL-6-mediated STAT3 phosphorylation in iTregs.** Histograms show the expression of pSTAT1 and pSTAT3 among Treg cells polarized *in vitro* in presence of TGF-β, IL-2 and IL-6 with or without IFN-γ (10 ng/mL, gate $CD4^+$ $Foxp3^+$).
