## Supplemental Fig 9 for "Multiple environmental signaling pathways control the differentiation of RORγt-expressing regulatory T cells"

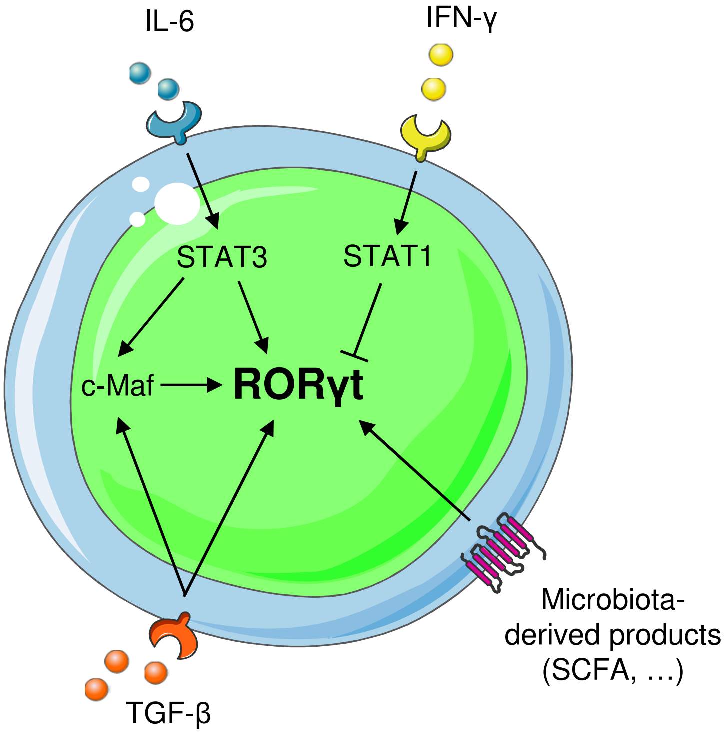


Figure S9. **Model of RORγt induction in Tregs.** The differentiation of specialized cell subsets is a balancing act: multiple positive and negative signals are integrated in order to tailor cell specialization to the immune context. Signals derived from a complex microbiota, or IL-6/STAT3 and TGF-β signaling induce RORγt expression in Tregs in a c-Maf-dependent or independent fashion. In return, inflammatory IFN-γ/STAT1 signaling opposes RORγt expression in Tregs in a c-Maf independent fashion.
